# Reciprocal interaction between I_K1_ and I_f_ in biological pacemakers: A simulation study

**DOI:** 10.1101/2020.07.23.217232

**Authors:** Yacong Li, Kuanquan Wang, Qince Li, Jules C. Hancox, Henggui Zhang

**Author notes:** Correspondence: Henggui Zhang.

## Abstract

Pacemaking dysfunction (PD) may result in heart rhythm disorders, syncope or even death. Current treatment of PD using implanted electronic pacemaker has some limitations, such as finite battery life and the risk of repeated surgery. As such, the biological pacemaker has been proposed as a potential alternative to the electronic pacemaker for PD treatment. Experimentally it has been shown that bio-engineered pacemaker cells can be generated from non-rhythmic ventricular myocytes (VMs) by knocking down genes related to the inward rectifier potassium channel current (I_K1_) or by overexpressing hyperpolarization-activated cyclic nucleotide gated channel genes responsible for the “funny” current (I_f_). Such approaches can turn the VM cells into rhythmic pacemaker cells. However, it is unclear if a bio-engineered pacemaker based on the modification of I_K1_- and I_f_-related channels simultaneously would enhance the ability and stability of bio-engineered pacemaking action potentials (APs). This study aimed to investigate by a computational approach the combined effects of modifying I_K1_ and I_f_ density on the initiation of pacemaking activity in human ventricular cell models. First, the possible mechanism(s) responsible for VMs to generate spontaneous pacemaking APs by changing the density of I_K1_ and I_f_ were investigated. Then the integral action of targeting both I_K1_ and I_f_ simultaneously on the pacemaking APs was analysed. Our results showed a reciprocal interaction between I_K1_ and I_f_ on generating stable and robust pacemaking APs in VMs. In addition, we thoroughly investigated the dynamical behaviours of automatic rhythms in VMs in the I_K1_ and I_f_ parameter space, providing optimal parameter ranges for a robust pacemaker cell. In conclusion, to the best of our knowledge, this study provides a novel theoretical basis for generating stable and robust pacemaker cells from non-pacemaking VMs, which may be helpful in designing engineered biological pacemakers for application purposes.

**Author Summary:** Pacemaking dysfunction has become one of the most serious cardiac diseases, which may result in arrhythmia and even death. The treatment of pacemaking dysfunction by electronic pacemaker has saved millions of people in the past fifty years. But not every patient can benefit from it because of possible limitations, such as surgical implication and lack of response to autonomic stimulus. The development of bio-pacemaker based on gene engineering technology provides a promising alternative to electronic pacemaker by manipulating the gene expression of cardiac cells. However, it is still unclear how a stable and robust bio-pacemaker can be generated. The present study aims to elucidate possible mechanisms responsible for a bio-engineered pacemaker by using a computational electrophysiological model of pacemaking cells based on modifying ion channel properties of I_K1_ and incorporating I_f_ in a human ventricular cell model, mimicking experimental approaches of gene engineering. Using the model, possible pacemaking mechanisms in non-pacemaking cells, as well as factors responsible for generating robust and stable biological pacemaker, were investigated. It was shown that the reciprocal interaction between reduction of I_K1_ and incorporation of I_f_ played an important role for producing robust and stable pacemaking. This study provides a novel insight into understanding of the initiation of pacemaking behaviours in non-rhythmic cardiac myocytes, providing a theoretical basis for experimental designing of biological pacemakers.

## Introduction

Currently, the electronic pacemaker implantation is the only non-pharmacological therapy for some patients with pacemaking dysfunctions (PD), such as sick sinus syndrome and atrioventricular heart-block, but it has some possible limitations (1). Implantation of pacemaker device may have complications for patients, especially for aged ones because of their infirm health (2). Pediatric patients can receive electronic pacemakers; however, the device has to be replaced as they grow and repeated surgeries are needed (3). Electronic devices can be subject to electromagnetic interference (4), which causes inconvenience to the patients. A further issue is that classical electronic pacemakers are insensitive not only to hormone stimulation (5) but also to autonomic emotion responsiveness (4), although there are some attempts to make them respond to autonomic nervous control (6). In addition, the long-term use of electronic pacemakers has been reported to increase the risk of heart failure (7). Appropriately designed biological pacemakers (bio-pacemakers) have the potential to overcome some of the limitations of electrical device use (8). For example, engineered bio-pacemakers could potentially involve only minor surgical trauma for implantation as well as opportune chronotropic responses to creature emotion (9). In previous experimental studies, it has been shown that a bio-pacemaker can be engineered *via* adenoviral gene transduction (10-12) or lentiviral vector (13, 14) techniques, by which non-pacemaking cardiac myocytes (CMs) can be transformed to rhythmic pacemaker-like cells.

The native cardiac primary pacemaker, sinoatrial node (SAN), is a special region comprised of cells with distinct electrophysiological properties to cells in the working myocardium. Such intrinsic and special electrophysiological properties of SAN cells are mainly manifested by their small if not absent inward rectifier potassium channel current (I_K1_) (15), but a large “funny” current (I_f_) (16) that is almost absent in atrial and ventricular cells. In addition to absence of I_K1_ and presence of I_f_, T-type Ca^2+^ channel current (I_CaT_) (17) and sustained inward current (I_st_) (18) also contribute to spontaneous pacemaking activity in SAN cells. Such unique electrophysiological properties of SAN cells form a theoretical basis to engineer non-pacemaking CMs into spontaneous pacemaker cells. These non-pacemaker cells include native CMs, such as ventricular (11, 19-21), atrial (22) or bundle branch myocytes (23). They can also be stem cells, such as embryonic stem cells (24-26), bone marrow stem cells (13, 27, 28), adipose-derived stem cells (29-31), or induced pluripotent stem cell (14, 32, 33).

With gene therapy, these cells are manipulated to provoke automaticity. In previous studies, knocking off the *Kir2*.*1* gene to reduce the expression of I_K1_ promoted spontaneous rhythms in newborn murine ventricular myocytes (VMs) (19); by reprogramming the *Kir2*.*1* gene in guinea-pigs, VMs also produced pacemaker activity when I_K1_ was suppressed by more than 80% (11, 20). As I_f_ plays an important role in the native SAN cell pacemaking, a parallel gene therapy manipulation to create engineered bio-pacemaker has been carried out by expressing the *HCN* gene family in non-rhythmic cardiac myocytes (34). It has been shown that expressing *HCN2* produced escape beats in canine CMs (22, 23) and initiated spontaneous beats in neonatal rat VMs (21). *HCN* expression in stem cells-induced-CMs also enhanced their pacemaking rate (13, 27, 28, 35, 36). Overexpressing *HCN4* can also induce spontaneous pacemaking activity in mouse embryonic stem cells (37). However, acute expression of the *HCN* gene might have a side effect on the normal cardiac pacemaking activity (38-40) and the overexpression of the *HCN* gene in VMs can cause ectopic pacemaker automaticity and even arrhythmicity (41).

It has been suggested that a combined manipulation of I_K1_ and I_f_ may be a better alternative for creating a bio-pacemaker (42). It has been demonstrated that the expression of transcriptional regulator *TBX18*, which influenced both I_f_ and I_K1_ expression, generated appropriate autonomic responses in non-pacemaking CMs (10, 31, 43). In addition, reprogramming *TBX18* in porcine VMs did not show the increase of arrhythmia risk (12), indicating the probable superiority of manipulating I_K1_ and I_f_ jointly for generating a bio-pacemaker. Furthermore, Chen et al. (44) attempted to explore the generation of oscillation by manipulating the expressing level of *Kir2*.*1* and the *HCN* genes and suggested that a dynamic balance between *Kir2*.*1* and *HCN* was essential to initiate oscillation in HEK293 cells. In the absence of I_K1_, spontaneous rhythmic oscillations might be inhibited due to an insufficiently repolarized membrane potential to activate I_f_. However, HEK293 cells lack all the other ionic channels present in native SAN myocytes.

Computational modelling offers a means to investigate different approaches to generating stable pacemaking activity. It has been used to study possible roles of down-regulating I_K1_ in VMs (45-47) and combined action of overexpression of I_f_ with down-regulation of I_K1_ in inducing spontaneous pacemaking activity in SAN (42). Bifurcation analysis has also been used to explore the effect of changes in some individual ion channel current on the pacemaking activities of SAN cells (48) and genesis of automaticity in VMs (46, 49), showing the role of I_K1_, I_f_, I_st_ and Na^+^/Ca^2+^ exchange current (I_NaCa_) in modulating the initiation and stability of pacemaking activity (49). But this approach was applied to simplified CMs model (46) and interplay between more than one ion channel currents on modulating bio-pacemaker APs has not been comprehensively investigated yet.

In this study, we constructed a bio-pacemaker model based on a human non-rhythmic VMs model (50) by manipulating I_K1_ and incorporating I_f_ (51) into the model. The aim of this study was to investigate (i) possible mechanism(s) underlying the pacemaking activity of the VMs in the I_K1_ and I_f_ parameter space; and (ii) the reciprocal interaction of reduced I_K1_ and increased I_f_ in generating stable pacemaking action potentials (APs) in them. In addition, possible factors responsible for impaired pacemaking activity due to inappropriate ratio between I_K1_ and I_f_ were also investigated. This study provides insightful understandings to generating stable and robust engineered bio-pacemaker.

## Results

### Initiation of transient spontaneous depolarization

In the basal VM cell model with the suppression of I_K1_ by 70% (the density of I_K1_ at −80 mV was 0.297 pA/pF), incorporating of I_f_ (with a current density of −0.63 pA/pF at −80 mV in the I-V curve of I_f_ (S1B Fig)) was unable to depolarize the membrane potential and lead to spontaneous pacemaking activity because the excessive outward current of I_K1_ counteracted the inward depolarizing current. This state can be described by State-1 as shown in Eq. 8. However, when the density of I_f_ was increased to −1.89 pA/pF, spontaneous depolarization was provoked at the beginning of the transition period, however, the automaticity self-terminated after 1.63×10^5^ ms (Fig 1 A), showing a State-2 behaviour as described in Eq. 9.

**Fig 1.**
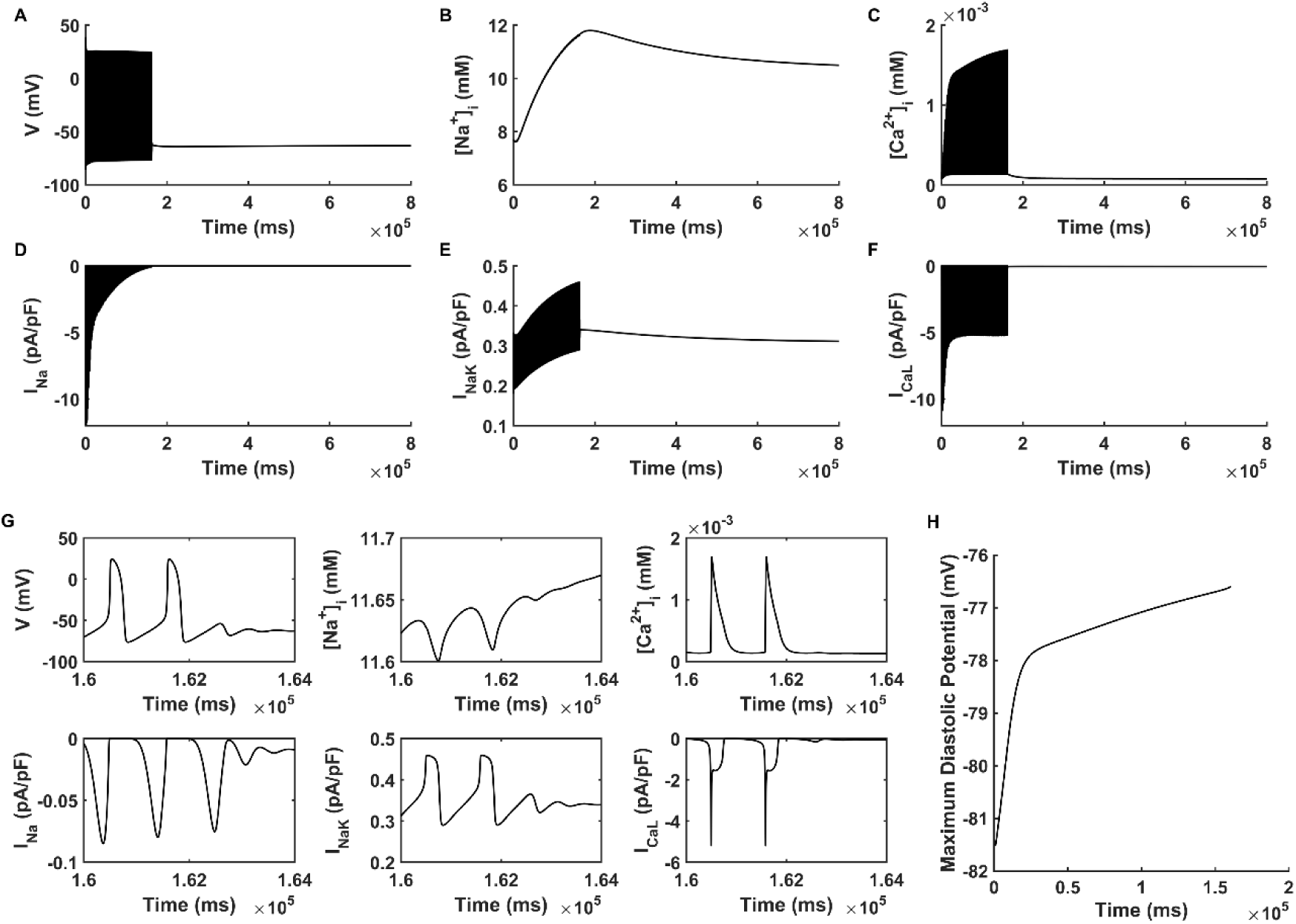
Transient spontaneous pacemaking behaviour. (A-F) Membrane potential (V), intracellular Na^+^ concentration ([Na^+^]_i_), intracellular Ca^2+^ concentration ([Ca^2+^]_i_), fast sodium current (I_Na_), L-type calcium channel current (I_CaL_) and Na^+^/K^+^ pumping current (I_NaK_) with the current densities of (I_K1_, I_f_) at (0.297pA/pF, −1.89 pA/pF) during the entire simulating period of 800 s. (G) Expanded plots of (A-F) for the time course of pacemaking self-termination (1.6× 10^5^ ms to 1.64× 10^5^ ms). H: Maximum diastolic potential of spontaneous pacemaking behaviour of (A).

We analysed possible ion channel mechanisms responsible for unstable and self-terminating pacemaking APs with the current densities of (I_K1_, I_f_) at (0.297pA/pF, −1.89 pA/pF) (‘CASE 1’). Results in Fig 1 showed that during the time course of the spontaneous pacemaking, there were changes of intracellular ionic concentrations and the MDP. Through the Na^+^ permeability of I_f_, there was extra Na^+^ flowing into the cytoplasm during each of the APs, leading the intracellular Na^+^ concentration ([Na^+^]_i_) to increase from 7.67 to 11.8 mM (Fig 1 B). The increased [Na^+^]_i_ augmented the feedback mechanism of Na^+^/K^+^ pumping activity, by which the Na^+^/K^+^ pump current (I_NaK_) increased gradually with time (Fig 1 E). In addition, there was also an accumulation of the intracellular Ca^2+^ concentration ([Ca^2+^]_i_, Fig 1 C) during the time course of spontaneous pacemaking APs. Such an accumulation of [Ca^2+^]_i_ was due to the fact that the automaticity in VMs shortened the DI between two successive APs, leaving insufficient time for Ca^2+^ in the cytoplasm to be extruded to restore to its initial value after each cycle of excitation. This consequentially led to overload in [Ca^2+^]_i_, which suppressed the extent of the activation degree of the L-type calcium current (I_CaL_, Fig 1 F), especially during the phase 0 of the pacemaking action potential. Furthermore, the overloaded [Ca^2+^]_i_ increased the I_NaCa_ (S2 Fig) gradually with time, resulting in an elevated MDP (Fig 1 H) that inhibited the activation degree of the fast sodium channel current (I_Na_, Fig 1 D). All of these factors worked together, inhibiting the membrane potential to reach the take-off potential, leading to self-terminated automaticity at 1.63×10^5^ ms (Fig 1 G).

It was also possible to generate automaticity in the model by fixing the current density of I_f_ at a low value, but with a further reduction in I_K1_ density. S3 Fig shows the results when I_f_ was held at −0.63 pA/pF, the current density of I_K1_ was reduced to 0.178 pA/pF. In this case, pacemaking activity appeared in the model, but the automaticity was unstable and self-terminated due to similar mechanisms as shown in Fig1 for the increased-I_f_ situation.

### Bursting pacemaking behaviour

Fig 2 shows the intermittent bursting behaviour, which is generated with a different combination of I_K1_ and I_f_ current densities in the model. In the figure, the current densities of (I_K1_, I_f_) were held at (0.297 pA/pF, −2.52 pA/pF) (defined as ‘CASE 2’). Such kind of pacemaking state can be classified as State-3 (Eq. 10).

**Fig 2.**
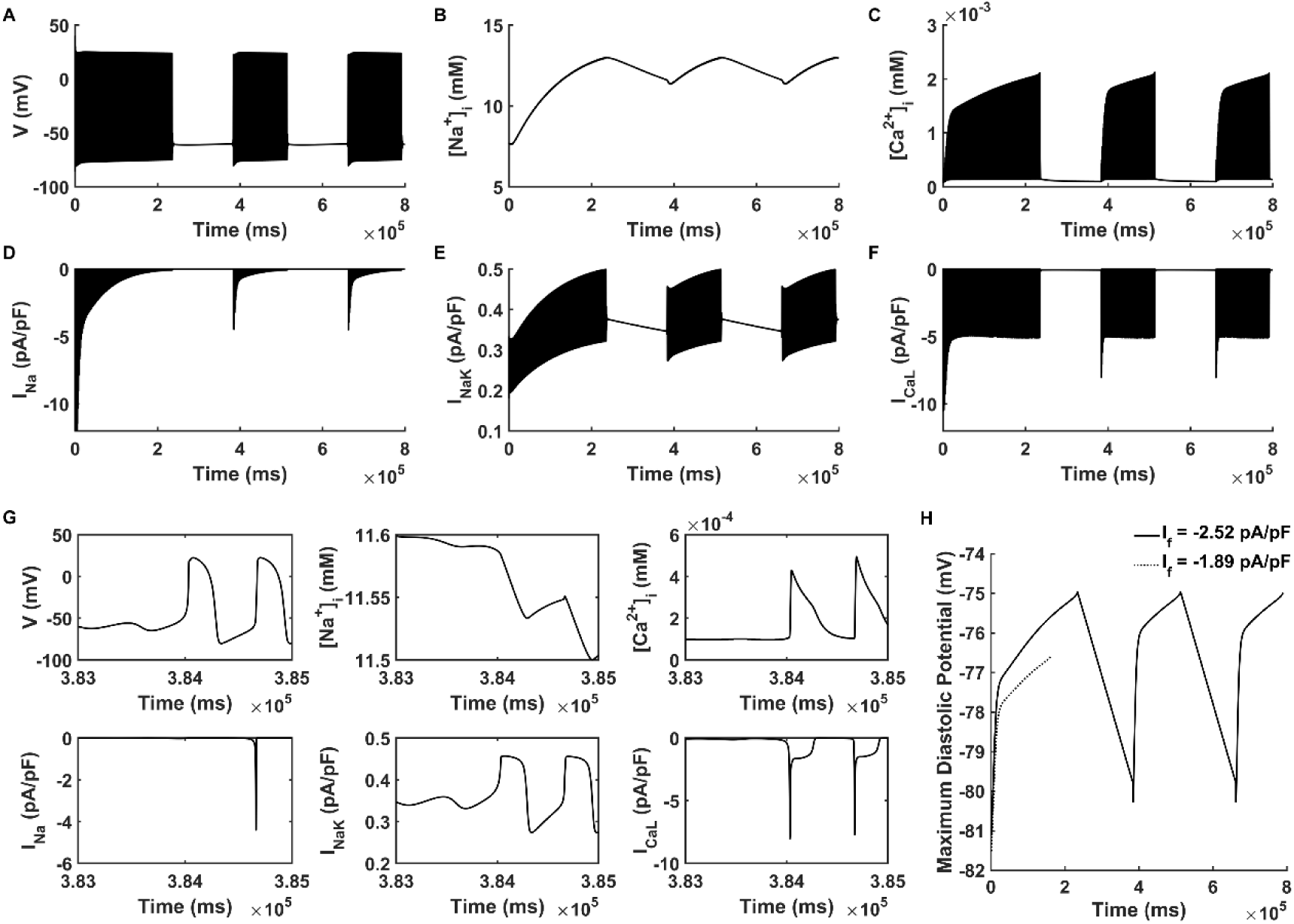
Bursting pacemaking behaviour. (A-F) Membrane potential (V), intracellular Na^+^ concentration ([Na^+^]_i_), intracellular Ca^2+^ concentration ([Ca^2+^]_i_), fast sodium channel current (I_Na_), Na^+^/K^+^ pumping current (I_NaK_) and L-type calcium channel current (I_CaL_) with the current densities of (I_K1_, I_f_) at (0.297 pA/pF, −2.52 pA/pF) during the entire simulating period of 800 s. (G) Expanded plots of (A-F) for the time course of pacemaking resumption (3.83×10^5^ ms to 3.85×10^5^ ms). H: Maximum diastolic potential of automatic pacemaking activity when I_f_ is −2.52 and −1.89 pA/pF (solid and dotted line respectively).

In CASE 2, the spontaneous oscillation was unstable, characterized by self-termination and then resumption after a quiescent period (Fig 2 A). Similar to CASE 1, the self-termination was accompanied by the overload of [Na^+^]_i_ (Fig 2 B), the accumulation of [Ca^2+^]_i_ (Fig 2 C), which caused the reduction of I_Na_ (Fig 2 D), the increase of I_NaK_ (Fig 2 E) and the decrease of I_CaL_ (Fig 2 G). This suggested that the underlying mechanisms responsible for the self-termination of the pacemaking APs were similar to those of CASE 1.

It is of interest to analyse the mechanism(s) for the resumption of the pacemaking APs after a long pause. It was shown that, during the time course of the quiescent interval (from 2.36×10^5^ ms to 3.84×10^5^ ms in Fig 2 A-F), the intracellular Na^+^ (Fig 2 B) continued to be extruded out of the cell by I_NaK_ (Fig 2 E), and the intracellular Ca^2+^ (Fig 2 C) extruded by the I_NaCa_ (S4A Fig). As such, the [Na^+^]_i_ (Fig 2 B) and [Ca^2+^]_i_ (Fig 2 C) gradually decreased over time. A decrease in [Na^+^]_i_ led to a gradually reduced I_NaK_ over the time course of quiescence (Fig 2 E), which decreased its suppressive effect on depolarization. During the time period of quiescence, I_CaL_ kept at a small magnitude (Fig 2 F). *Via* I_NaCa_ (S4 Fig), the intracellular Ca^2+^ was kept to be extruded out of the cell, leading to a decreased [Ca^2+^]_i_. Consequentially, a decrease in [Ca^2+^]_i_ resulted in a reduced calcium-dependent inactivation of I_CaL_, leading to an increased I_CaL_ (Fig 2 G), which facilitated the action potential generation. Moreover, as compared with CASE 1, the increase in I_f_ also helped to produce a more depolarized MDP (Fig 2 H), allowing the membrane potential more easily to reach the take-off potential for initiation of the upstroke. All of these actions worked in combination to produce a full course of action potential with a sufficient amplitude that activated sufficient outward currents to repolarize the cell membrane to a range that activated I_f_ and I_Na_, facilitating the resumption of the spontaneous pacemaking activity at 3.847×10^5^ ms (Fig 2 G). This process of self-termination and resumption repeated alternately, which constituted bursting behaviour.

### Persistent pacemaking activity

A further increase in I_f_ ((I_K1_, I_f_) at (0.297 pA/pF, −3.15 pA/pF)) produced a series of persistent spontaneous APs. Results are shown in Fig 3 (grey lines) for APs (Fig 3 A), together with a phase portrait of membrane potential (V) and total membrane channel current (I_total_) (Fig 3 B), I_K1_ (Fig 3 C), and I_CaL_ (Fig 3 D). Although the spontaneous APs were sustained during the entire simulation period of 800 s, there were some incomplete depolarizations observed periodically (Fig 3 A, grey line) which can be classified as State-4 according to Eq. 11. This pacemaking situation was termed as ‘CASE 3’.

**Fig 3.**
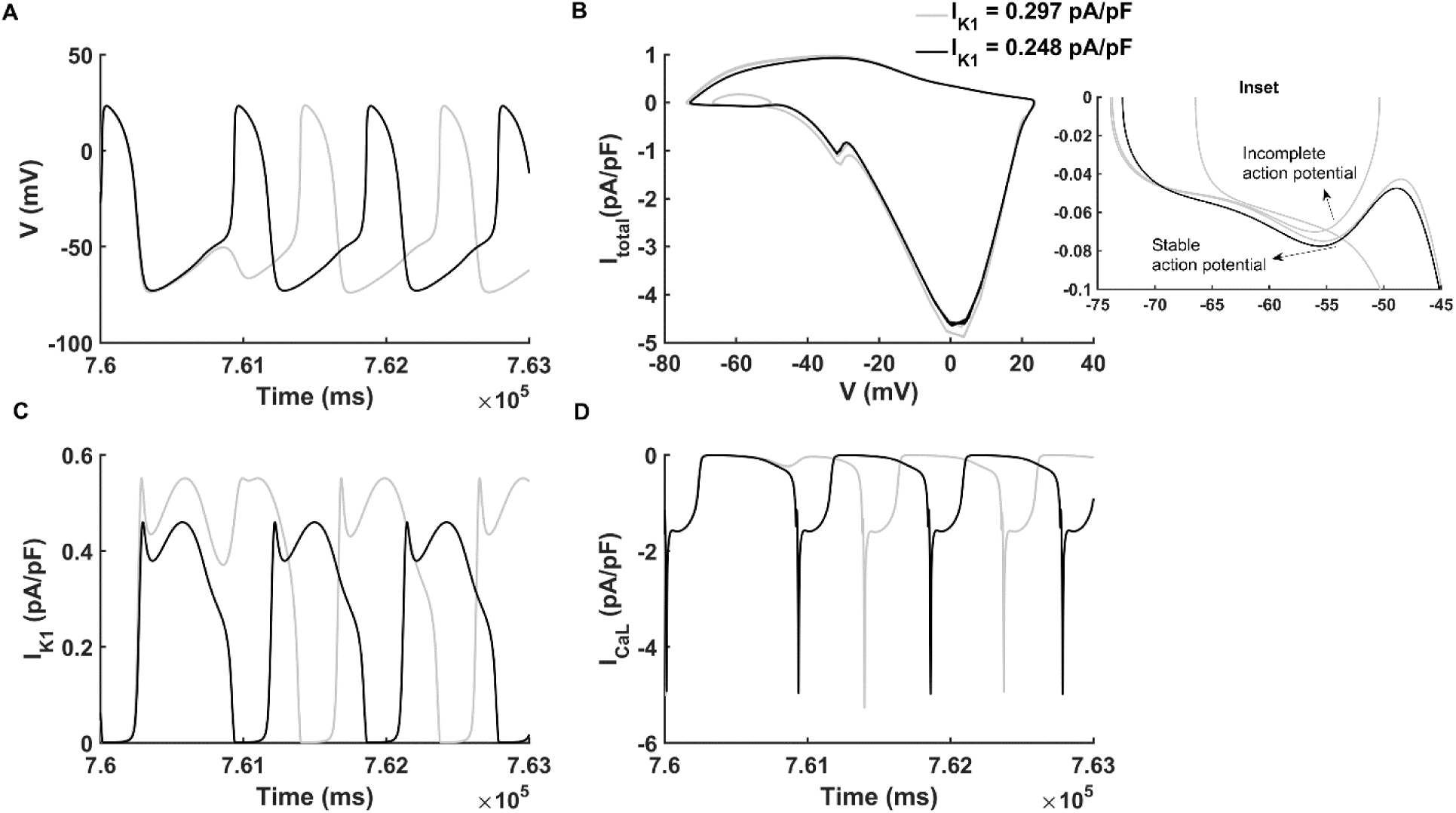
Persistent pacemaking activity. The current densities of (I_K1_, I_f_) of stable pacemaking activity and periodically incomplete pacemaking activity are at (0.248 pA/pF, −3.15 pA/pF) (black lines) and (0.297 pA/pF, −3.15 pA/pF) (grey lines) respectively. (A-D) The membrane potential (V), phase portraits of membrane potential against the total membrane channel current (I_total_), inward rectifier potassium channel current (I_K1_) and L-type calcium channel current (I_CaL_) with simulating time course from 7.6× 10^5^ to 7.63× 10^5^ ms. (Inset) Expanded plot for phase diagram during V from −75 to −45 mV and I_total_ from −0.1 to 0 pA/pF.

When the density of I_K1_ was further reduced to 0.248 pA/pF (Fig 3 C, black line), a stable pacemaking activity was established (Fig 3A, black line), with an average CL of 895 ms and MDP of −72.63 mV. This kind of pacemaking state can be described as State-5 by Eq. 12. In this condition, the pacemaking activity was robust and the pacing CL was close to that of the native human SAN cells (approximately 800 - 1000 ms (52)). We termed stable pacemaking activity with (I_K1_, I_f_) at (0.248 pA/pF, −3.15 pA/pF) as ‘CASE 4’.

In order to understand potential mechanism(s) underlying the genesis of incomplete pacemaking potentials in CASE 3, phase portraits of membrane potential against I_total_ for incomplete (grey line) and complete (black line) depolarization APs were plotted and superimposed for comparison, as shown in Fig 3 B, with a highlight of phase portraits during the diastolic pacemaking potential range from −75 mV to about −45 mV being shown in the inset. In the case of incomplete depolarization (CASE 3), there was a greater I_K1_ (Fig 3 C) that counteracted the inward depolarizing current, leading to a smaller I_total_ during the diastolic depolarization phase (see the grey line in Fig 3 B and the inset). Consequentially the membrane potential failed to reach the take-off potential for the activation of I_CaL_ (Fig 3 D, grey line), leading to an incomplete course of action potential (Fig 3 A, grey line). In CASE 4, with a reduced I_K1_ (Fig 3 C, black line), there was a greater I_total_ during the diastolic depolarization phase (Fig 3 B and the inset, black line), which drove the membrane potential to reach the take-off potential for the activation of I_CaL_ (Fig 3 D, black line), leading to the upstroke of the action potential.

The frequency for the appearance of the incomplete AP was dependent on the density of I_K1_. Incomplete depolarization occurred less frequently, with progressively smaller I_K1_. By way of illustration, the incomplete depolarization appeared once every three cycles with the I_K1_ density at 0.297 pA/pF in CASE 3 (S5A Fig), but this became once every five cycles when I_K1_ density was reduced to 0. 277 pA/pF I_K1_ (S5B Fig). This suggested that a large residual of I_K1_ in the VM model might result in the failure of complete depolarization.

### Dynamic analysis in I_K1_ and I_f_ parameter space

Dynamic pacemaker AP behaviours were dependent on the balance of I_K1_ and I_f_ interactions. Further simulations were conducted in the I_K1_ and I_f_ parameter space to characterize this dependence. Results are shown in Fig 4. With differing combinations of I_K1_ and I_f_ density, five different regions for distinctive pacemaking dynamics could be discerned, including stable pacemaking activity (blue area, features described by Eq. 12), intermittence of failed depolarization (yellow area, features described by Eq. 11), bursting pacemaking behaviour (orange area, features described by Eq. 10), transient pacemaking activity (green area, features described by Eq. 9), and no automaticity (grey area, features described by Eq. 8). In each category, representative membrane potentials are illustrated at the bottom panel of the Fig 4.

**Fig 4.**
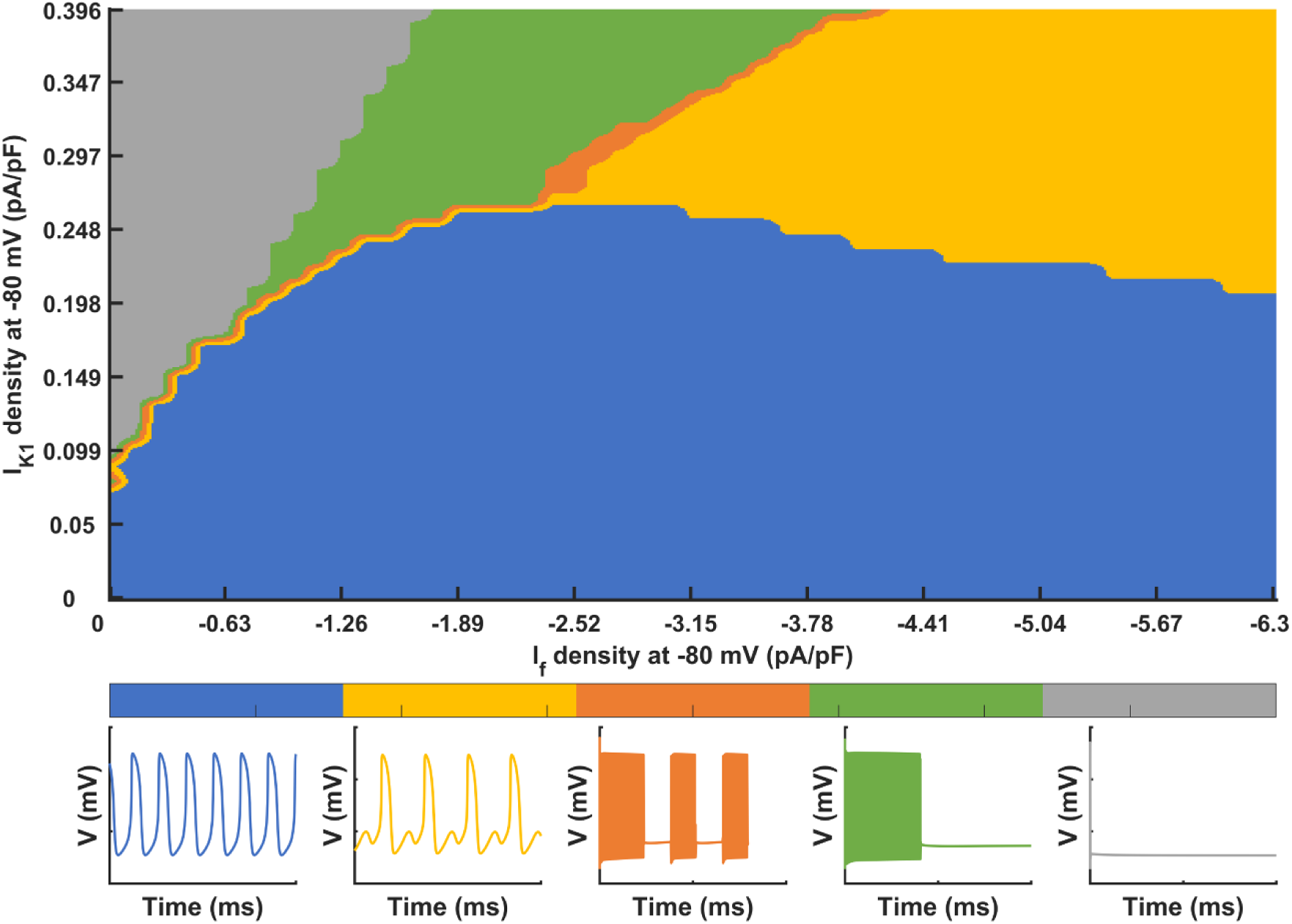
The dynamical behaviours of the pacemaking action potentials in I_K1_ and I_f_ parameter space. Blue: stable pacemaking activity; Yellow: persistent pacemaking activity with periodic incomplete depolarization. Orange: bursting pacemaking behaviour. Green: transient spontaneous pacemaking behaviour. Gray: no spontaneous pacemaking behaviour. In each category, the typical pacemaking action potentials are illustrated at the bottom panel.

With a fixed I_f_ density, alterations to I_K1_ could produce different types of pacemaking activities and this also applied when I_K1_ was fixed whilst I_f_ was changed. To illustrate, when I_f_ density was fixed at a density between −0.63 and −2.52 pA/pF, with a 60 – 80% block of I_K1_ (i.e., I_K1_ density at −80 mV was in the range of 0.198-0.396 pA/pF), pacemaking activity was generated but with self-termination (Fig 4, green area). Then, a further reduction in I_K1_ or a slight increase in I_f_ induced bursting pacemaking behaviour, as shown by the orange area in Fig 4, which was between the boundaries marking the persistent automaticity and transient pacemaking activity regions. A further increase in I_f_ or suppression in I_K1_ could produce persistent automaticity (Fig 4, yellow and blue area). But when I_K1_ was greater than about 0.248 pA/pF, incomplete depolarization appeared periodically (Fig 4, yellow area). Finally, a stable and spontaneous pacemaking activity could be generated when I_K1_ was decreased to less than 0.248 pA/pF at −80 mV with I_f_ included (Fig 4, blue area).

### Reciprocal role of I_f_ and I_K1_ in generating pacemaking APs

Further analysis was conducted to investigate the reciprocal role of reduced I_K1_ and increased I_f_ in generating pacemaking APs. By sufficiently reducing I_K1_ to a density of 0.05 pA/pF alone, the model was able to generate spontaneous APs with a CL of 1011 ms. In this case, the incorporation of I_f_ with a small density helped to boost the pacemaking activity and increase the pacemaking frequency. It was shown that with the incorporation of I_f_ at a density of 0.63 pA/pF, the CL was reduced by 233 ms, changing from 1011 ms to 778 ms (Fig 5 A). Compared with the case of I_f_ absence, incorporation of I_f_ - even with a small density (−0.63 pA/pF) - helped to depolarize cell membrane potential during the early DI phase (Fig 5 A and C). Moreover, the incorporation of I_f_ led to an accumulation of [Na^+^]_i_ (Fig 5 B) then increased amplitude of I_NaCa_ (Fig 5 D), which also contributed to the depolarization of membrane potential. As a result, the incorporation of I_f_ facilitated genesis of spontaneous APs and shortened the DI significantly, thus decreased the CL.

**Fig 5.**
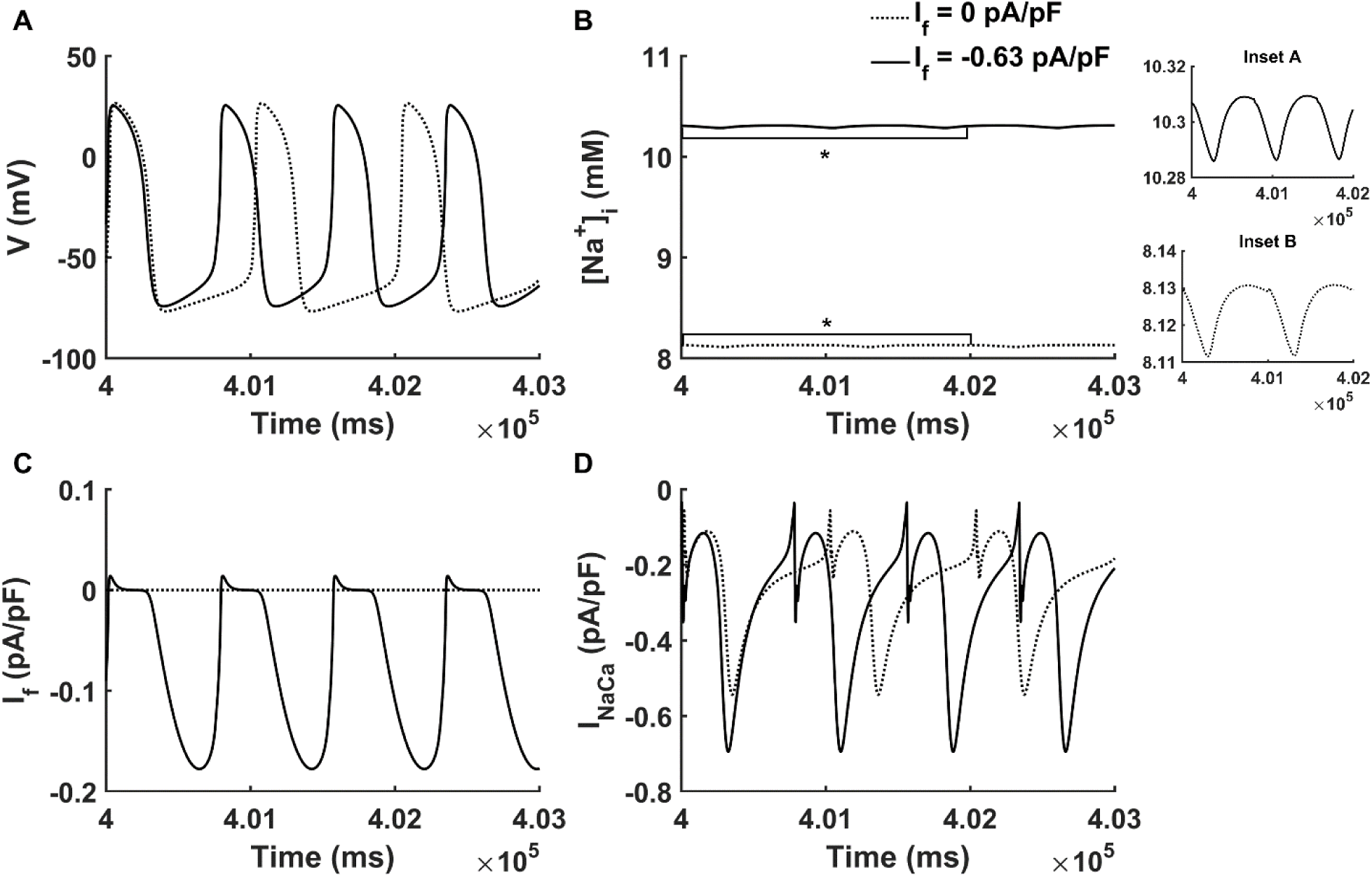
Role of I_f_ in pacemaking ability. (A-D) The membrane potential (V), intracellular Na^+^ concentration ([Na^+^]_i_), hyperpolarization-activated channel current (I_f_), Na^+^/Ca^2+^ exchange current (I_NaCa_) with simulating time course from 4×10^5^ to 4.03×10^5^ ms when the current densities of (I_K1_, I_f_) are at (0.05 pA/pF, 0 pA/pF) and (0.05 pA/pF, −0.63 pA/pF) (dotted and solid line respectively). (Inset A-B) Expanded plots of [Na^+^]_i_ traces for the time course marked by the horizontal brackets with asterisks in (B).

With a fixed I_K1_ density of 0.05 pA/pF, the relationship between the computed CL of pacemaking APs and I_f_ density was found to be nonlinear as shown in Figure 6 A. In a range from −0.126 to −2.52 pA/pF, an increase in I_f_ density produced a marked decrease in the CL (Fig 6 A), which was associated with an increase in the rate of membrane depolarization during the DI (diastolic depolarizing rate) (S6B Fig). In this range, an increase in I_f_ caused an elevated MDP (Fig 6 B), as well as an accumulation of [Na^+^]_i_ (Fig 6 C), which enhanced I_NaCa_ (Fig 6 D). All of these contributed to the acceleration of the pacemaking activity. However, when I_f_ density was over −2.52 pA/pF, there was a less dramatic decrease in CL with an increase of I_f_ (Fig 6 A). This was attributable to a reduced I_Na_ (Fig 6 F) as a consequence of gradual elevation of the MDP (Fig 6 B). Another factor was that the maximum density of I_f_ was not increased linearly (Fig 6 E) because of elevated MDP.

**Fig 6.**
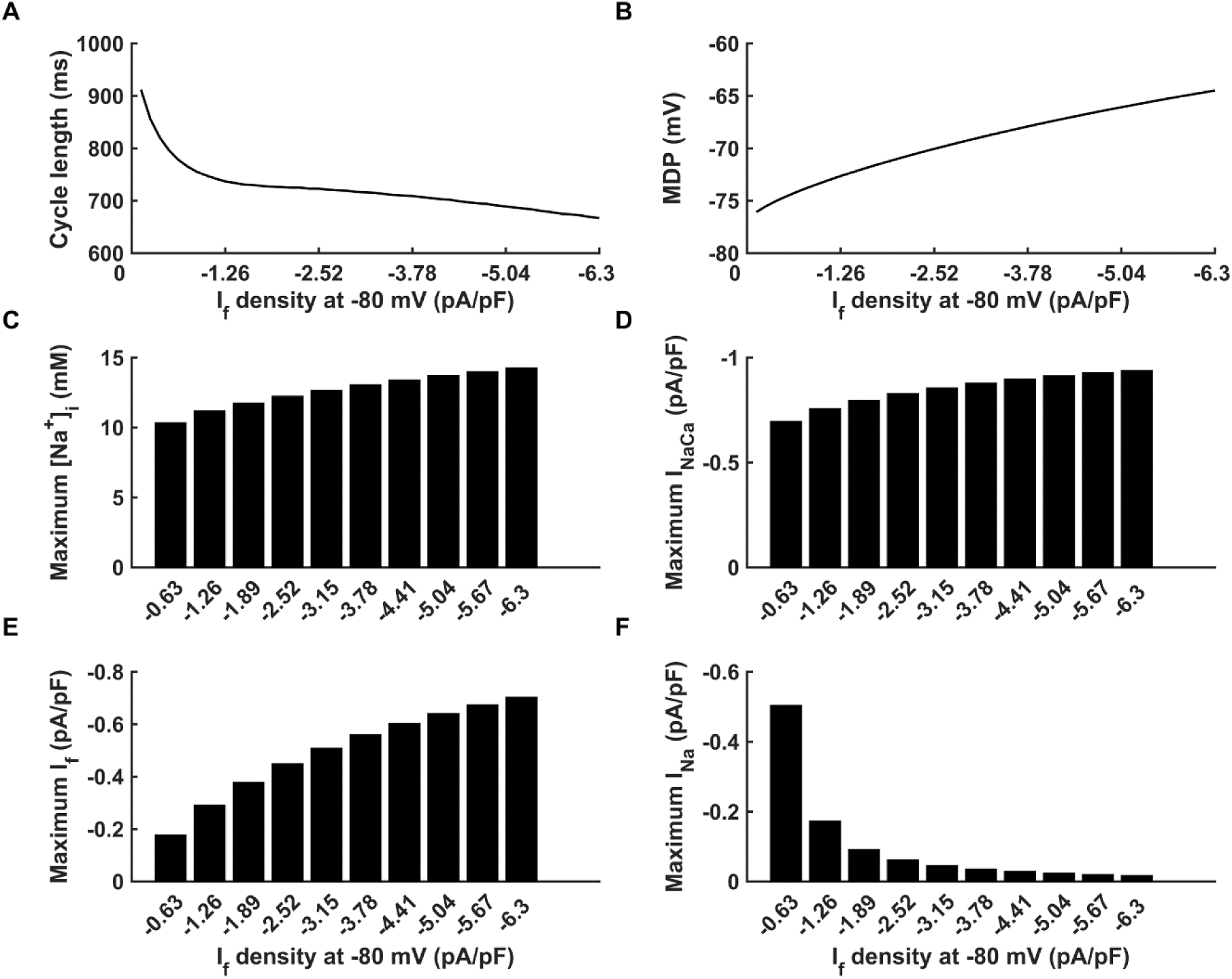
Effect of I_f_ density on the pacemaking cycle length under fixed I_K1_ density. I_K1_ density is fixed at 0.05 pA/pF. (A-B) Change of cycle length and maximum diastolic potential (MDP) with the increase of I_f_ from 0 to −6.3 pA/pF. (C-F) Change of maximum intracellular Na^+^ concentration ([Na^+^]_i_), maximum Na^+^/Ca^2+^ exchange current (INaCa), maximum funny current (I_f_) and maximum fast sodium current (I_Na_) during a pacemaking period with the increase of I_f_.

Depending on the I_K1_ density, the relationship between CL and I_f_ density could also be biphasic as shown in Fig 7. With a small I_K1_ density (0.05 pA/pF), the measured CL decreased monotonically with the increase in I_f_ density (Fig 7 A, dotted line). However, with a large I_K1_ (0.198 pA/pF), the measured CL first decreased with an increased I_f_ density, but then increased with it when I_f_ density was greater than −3.15 pA/pF, implicating a slowdown in the pacemaking activity with the increase of I_f_ (Fig 7 A, solid line). Such a slowdown of pacemaking APs with an increased I_f_ was mainly due to the prolonged DI (S7 Fig). This was attributable to the delicate balance between the total integral of three dominant inward currents (I_Na_, I_NaCa_ and I_f_) (I_in_ as defined in Eq. 14) and the total integral of four dominant outward currents (I_K1_, I_NaK_, I_Kr_ and I_Ks_) (I_out_ as defined in Eq. 15) during the DI phase (Fig 7 B and C). At a low I_K1_ density of 0.05 pA/pF, with the increase in I_f_ density, there was a monotonic increase in both of I_in_ and I_out_, and the balance of them resulted in a monotonic decrease of CL (Fig 7, dotted lines). However, at a large I_K1_ density (e.g. 0.198 pA/pF), an increase of I_f_ was associated with a monotonic increase in I_out_ (Fig 7 C, black line), but had a different impact on I_in_, which reached at a flat magnitude after I_f_ density was greater than −3.15 pA/pF (Fig 7 B, black line). Consequentially, at large I_f_, a greater I_out_ out balanced the I_in_, leading to a prolonged DI that increased the CL.

**Fig 7.**
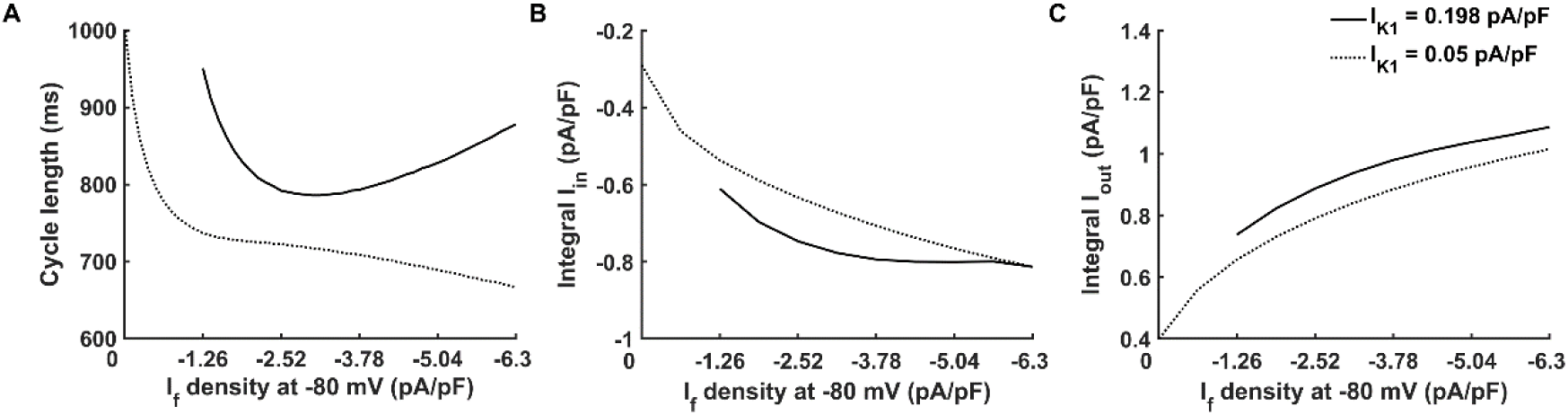
Effect of I_f_ density on the pacemaking cycle length under different I_K1_ density. I_K1_ density is 0.198 (solid line) and 0.05 pA/pF (dotted line) at −80 mV. (A) Change of cycle length with the increase of I_f_ from 0 to −6.3 pA/pF. (B) Change of the total integral of main inward currents (Integral I_in_) during diastolic interval phase. The inward currents include fast sodium current (I_Na_), Na^+^/Ca^2+^ exchange current (I_NaCa_) and funny current (I_f_). (C) Change of the total integral of main outward currents (Integral I_out_) during diastolic interval phase. The outward currents include inward rectifier potassium channel current (I_K1_), Na^+^/K^+^ pumping current (I_NaK_), rapid delayed rectifier potassium channel current (I_Kr_) and slow delayed rectifier potassium channel current (I_Ks_).

### Pacemaking cycle length in I_K1_ and I_f_ parameter space

A systematic analysis of the relationship between the calculated CL in the I_K1_ and I_f_ density parameter space is presented in Fig 8. In the figure, the measured CL was coloured from 650 ms in dark red to 1000 ms in yellow. In this study, we regarded persistent pacemaking action potential with CL 1000 ms or less as ‘valid pacemaking activity’, therefore, only the CLs of the valid pacemaking potentials are shown in Fig 8.

**Fig 8.**
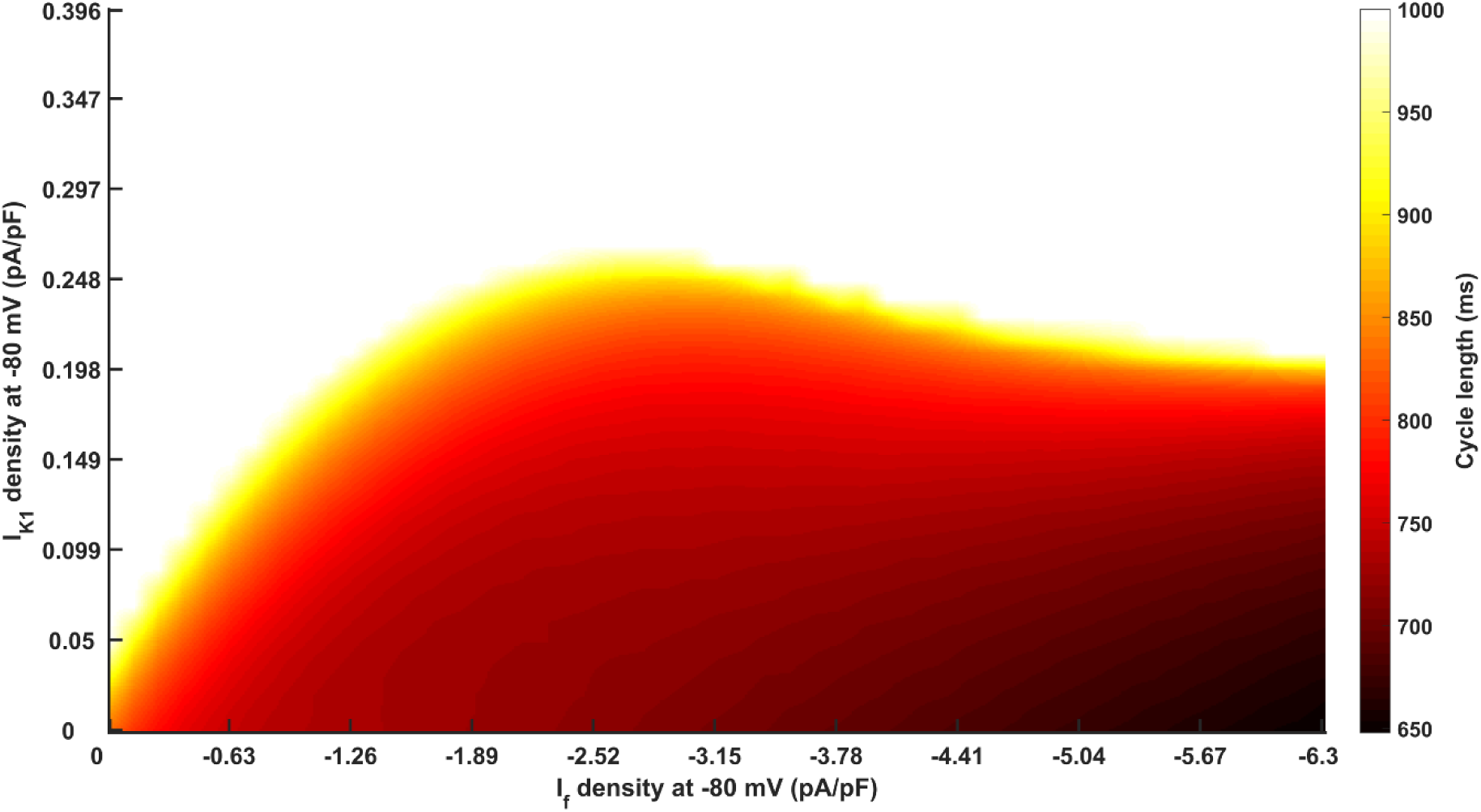
Measured cycle length in the I_K1_ and I_f_ density parameter space. The density of I_K1_ is from 0 to 0.396 pA/pF and the density of I_f_ is from 0 to −6.3 pA/pF at −80 mV. The measured CL is coloured from 650 ms in dark red to 1000 ms in yellow. White means that pacemaking cycle length is ‘invalid’ (more than 1000 ms) or membrane potential is not persistent during the whole simulating time course.

It was shown that a sufficient depression in I_K1_ (up to 75%; I_K1_ density < 0.248 pA/pF) was required to produce a stable pacemaking action potential with a ‘valid’ pacemaking frequency. With the increase of the I_K1_ inhibition level, the CL became shortened at all I_f_ densities considered. Also, the more that I_K1_ was inhibited, the less I_f_ was needed to provoke ‘valid’ spontaneous pacemaking activity. By contrast, the effect of I_f_ on the CL presented two phases, which was dependent on the I_K1_ density. When I_K1_ was less than 0.198 pA/pF, the pacemaking ability became robust with the increase in I_f_ density. However, when I_K1_ was increased from 0.198 to 0.248 pA/pF, an increase in I_f_ actually slowed the pacemaking activity, leading to an increased CL.

## Discussion

### Summary of major findings

In this study, we construct a virtual bio-engineered pacemaking cell model based on a human VM model by a combination of reduction of I_K1_ and incorporation of I_f_. Using the developed bio-pacemaker model, we investigate the combined actions of different I_K1_ and I_f_ permutations on the dynamical behaviours of membrane potential, ionic channel currents and intracellular ionic concentrations. It is shown that robust and stable pacemaking activity can be established by balancing the actions of reduced I_K1_ and increased I_f_, though the effect of each manipulation on pacemaking activity is different. While the action of a reduced I_K1_ on the pacemaking activity is consistent, that of an increased I_f_ is biphasic. Whilst the incorporation of I_f_ at an appropriate level promotes pacemaking activity, excessive I_f_ might result in abnormal pacemaking activity accompanied by abnormal intracellular ionic concentrations, which could be proarrhythmic. As a result, the reciprocal interaction between I_K1_ and I_f_ is crucial for producing stable spontaneous pacemaking activity in VMs. The results of this study may be useful for optimizing the future design of engineered bio-pacemakers.

### Role of I_K1_ suppression on pacemaking activity

The suppression of I_K1_ plays an important role in the generation of pacemaking activities in the VM cell model. Our simulation results have shown that a significant I_K1_ suppression (at least 60%) with the incorporation of I_f_ is required to provoke auto-rhythmicity in the model. With a modest suppression (i.e. 60-75% suppression of I_K1_), only unstable spontaneous APs can be produced. When I_K1_ is further suppressed by 75% - 100%, persistent, steady pacemaking behaviour can be initiated in our model with appropriate incorporation of I_f_ (Fig 4). These simulation results are in consistence with those of experimental findings, where it has been found that I_K1_ could be suppressed by 50 – 90 % by knocking out the Kir2.1 gene, and more than 80% inhibition of I_K1_ was required to produce a pacemaking phenomenon in guinea-pig’s ventricular cavity (11, 20). It is also in agreement with previous bifurcation analyses in showing that it required I_K1_ to be reduced to at least 15% of the control value to transform a VM cell model to be auto-rhythmic (46). and a complete block of I_K1_ produced a spontaneous pacemaking activity with a CL of 795 ms (46), close to 833 ms when I_K1_ was totally suppressed in the present study. We have also found a monotonic relationship between the measured CL and the degree of I_K1_ suppression. It suggests that when I_K1_ is inhibited enough to induce automaticity, the more the I_K1_ is blocked, the faster the pacemaking activity is with all I_f_ densities considered (Fig 8). Similar results have also been observed in another ventricular cell model developed for human VMs (53) based on modifications of the model of O’Hara and Rudy (54) (S1 Text and S8 Fig, solid and dotted lines).

Though our simulation results suggest an important role of sufficient suppression of I_K1_ for generating persistent and stable pacemaking APs, it is noteworthy that the deficiency of I_K1_ has been reported to be lethal for adult rodents (55); and loss function of *Kir2* gene may prolong QT intervals as well as cause Andersen’s syndrome (56). Consequently, suppression of I_K1_ from VM for generating a biological pacemaker may only be suitable when applied to highly localized, designated ‘pacemaker’ regions.

### Role of I_f_ in pacemaking activity

I_f_ has been shown to play an important role in generating pacemaking APs in both native (13, 21, 27, 28, 35, 36) and engineered pacemakers (42). Experimentally it has been shown that high expression of *HCN2* can initiate spontaneous beats in neonatal rat VMs (21, 35) and improve spontaneous beats in rabbit CMs (13). *HCN4* incorporation by the expression of *TBX18* can also initiate spontaneous pacemaking activity in both rodent VMs (10) and porcine VMs (12).

In the present study, we have also highlighted the role of I_f_ in generating pacemaking activity. Our simulation results have shown that with different I_f_ densities, the pacemaking activity may present different behaviours, including transient automaticity with self-termination, bursting behaviour, and persistent pacemaking (Fig 4). The incorporation of low amplitude of I_f_ can help to boost the pacemaking activity in the VM-based pacemaker model induced by I_K1_-inhibition (Fig 5). It helped to promote pacemaking activity, *via* its action of depolarization during the diastolic depolarization phase as well as its action on the intracellular ion concentrations. It has been shown that the inclusion of I_f_ in the VM cell model causes the accumulation of [Na^+^]_i_, which enhances Na^+^/Ca^2+^ exchange, thus promoting membrane potential depolarization especially during the early stage of DI (Fig 5). Such a promoting action of I_f_ in bio-pacemaking was also shown in another ventricular cell model as shown in S1 Text (S8 Fig, solid and dashed lines).

An increase in I_f_ density can enhance the automaticity in most cases. However, the effect of I_f_ on the pacemaking activity was observed to be biphasic. In our simulation, increasing I_f_ from a small initial density was associated with an increased pacemaking rate manifested by a decreased CL. But when it was increased to be over −2.52 pA/pF, excessive I_f_ resulted in an elevated MDP (Fig 6 B), which caused a reduced activation of I_f_ and I_Na_ (Fig 6 E and F), leading to a slowdown of the ability of pacemaking activity. The negative effect of excessive I_f_ on pacemaking APs was also observed in another ventricular pacemaker model (53) (S1 Text). A further increase in I_f_ density even terminated pacemaking activity (S9 Fig). This mechanism is verified by the fact that in the bio-pacemaker induced by HCN2 expression (57), co-expression of the skeletal muscle sodium channel 1 (*SkM1*), in order to hyperpolarize the action potential threshold, helped to counterbalance the negative effect of I_f_ overexpression, producing an accelerated depolarization phase. Furthermore, when I_K1_ was at a high value (e.g. density at 0.198 pA/pF), acute I_f_ even lengthened pacemaking period (Fig 7 A, black line). This simulation result is in agreement with a previous biological experimental study that observed a negative action of acute HCN gene expression in cardiac automaticity (41).

### Reciprocal interaction between I_K1_ and I_f_

Our study demonstrated that the reciprocal interaction between I_K1_ and I_f_ plays a crucial role in creating stable and persistent pacemaking. The present study has shown that only an optimal combination of I_K1_ and I_f_ can initiate stable pacemaking activity (Fig 4). In the presence of I_f_, the greater the degree of I_K1_ suppression, the smaller was the I_f_ density required for the generation of spontaneous oscillation (Fig 4). And modulation of the two currents simultaneously helps to create a physiologically-like pacemaker that is better than that produced by manipulating I_K1_ or I_f_ alone (Fig 8). Such observation of reciprocal interaction between I_K1_ and I_f_ in pacemaking is consistent with previous experimental observations. Previous studies have shown that although suppressing I_K1_ (11, 20), or incorporating sufficient I_f_ (21) alone was able to initiate pacemaking activity in VM cells, a pacemaker constructed by *TBX18* showed greater stability, due to its combined actions of I_K1_ reduction and I_f_ increase (10). Another experiment in porcine VMs (12) also indicated that *TBX18* expression did not increase the risk of arrhythmia, which means that a mixed-current approach is probably a superior means of producing a bio-pacemaker. Experiments in a *Kir2*.*1*/*HCN2* HEK293 cell (44) and *Kir2*.*1*/*HCN4* (42) showed that I_K1_ may actually recruit more I_f_ by activating current at more negative membrane voltages, because I_K1_ was the only negative current in these experiments. Our simulation, however, did not yield such a result because the interaction of other positive currents (such as I_NaK_, I_Kr_ and I_Ks_) contributed to the hyperpolarization of membrane potential and helped the activation of I_f_.

In addition, simulation results showed that I_K1_ expression level may influence the I_f_’s effect on the pacemaking activity. Excessive I_K1_ hindered I_f_’s ability to modulate pacemaking activity (Fig 7A and Fig 8). An experiment showed a coincident result that the expression of HCN2 in adult rat VMs could not cause spontaneous beats due to the high expression of I_K1_ (21), but in neonatal rats, the I_K1_ was less so that expressing HCN2 could provoke automaticity.

### Limitations

The limitations of human VMs model we used in this study has been described elsewhere (58). In this study, the I_f_ formulation of human SAN (51) was incorporated into the original VMs model. The properties of I_f_, including the conductance of I_f_, the half-maximal activation voltage (V_1/2_) and time constants of the activation, may present species-dependence. In this study, we only consider the conductance of I_f_ but have not discussed other properties of I_f_.

In addition, in this study, we only investigated the pacemaking action potential at the single-cell level, without considering the intercellular electrical coupling between pacemaker cells as presented in the SAN tissue. Mathematical analysis showed that the incorporation of I_st_ and I_CaT_ into VMs may promote the pacemaking ability of ventricular pacemaker in the coupled-cell model (49). However, up to now, there is no experimental study conducted yet to produce pacemaker cells from VMs by modifying the expression of I_st_ and I_CaT_, thus we did not discuss them in the present study. It is necessary to highlight these limitations, they nevertheless do not affect our conclusions on the underlying pacemaking mechanisms of engineered bio-pacemaker cells, especially regarding the reciprocal interaction of I_K1_ and I_f_ for a robust bio-pacemaker in modified VMs.

## Methods

### Single bio-pacemaker cell model

Previous experimental studies (10, 11, 19-21) implemented the suppression of *Kir2*.*1*, the incorporation of *HCN* channels and the expression of *TBX18* to induce pacemaking in VMs. In this study, we used a mathematical model of human VMs (50) as the basal model to investigate possible pacemaking mechanisms in VM-transformed pacemaking cells. In brief, the basal VM cell model can be described by the following ordinary differential equation:

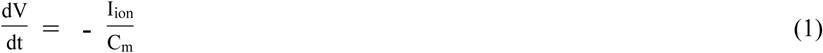

where V is the voltage across cell membrane surfaces, t is time, I_ion_ is the sum of all transmembrane ionic currents, and C_m_ cell capacitance.

The I_ion_ in the original ventricular model is described by the following equation:

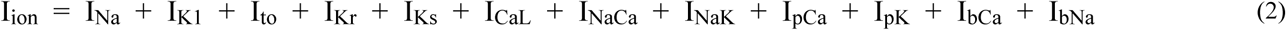

where I_Na_ is fast sodium channel current, I_K1_ is inward rectifier potassium channel current, I_to_ is transient outward current, I_Kr_ is rapid delayed rectifier potassium channel current, I_Ks_ is slow delayed rectifier potassium channel current, I_NaCa_ is Na^+^/Ca^2+^ exchange current, I_NaK_ is Na^+^/K^+^ pump current, I_pCa_ and I_pK_ are plateau Ca^2+^ and K^+^ currents, and I_bCa_ and I_bNa_ are background Ca^2+^ and Na^+^ currents. The formulations and their parameters for the ionic channels of human VM cells were listed in Ref. (50, 58).

To mimic the reduction of *Kir2*.*1* expression (11, 19, 20) or suppressing I_K1_ by expressing *TBX18* (10), in simulations, I_K1_ was decreased by modulating its macroscopic channel conductance (G_K1_). To mimic the incorporation of I_f_ in VMs experimentally (21), we modified the basal model of Eq. 2 by incorporating human SAN I_f_ formulation (51). In simulations, I_f_ was modulated by changing its channel conductance (G_f_).

As a result, the I_ion_ for the bio-pacemaker model can be described as:

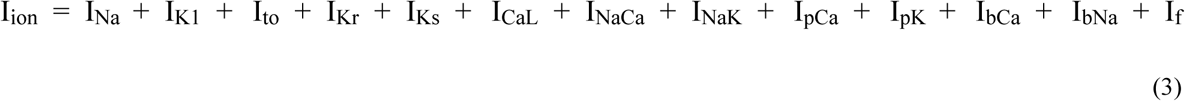

where I_K1_ could be expressed by

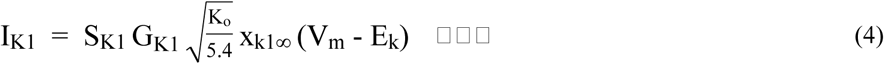

where G_K1_, *x*_*k*1∞_ is the conductance of I_K1_, is a time-independent inward rectification factor, K_o_ is extracellular K^+^ concentration and E_K_ is the equilibrium potentials of K^+^. S_K1_ is a scaling factor used to simulate the change of I_K1_ expression level.

I_f_ has two ionic channels and could permeate Na^+^ and K^+^ respectively. I_f_ could be described by

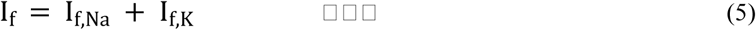

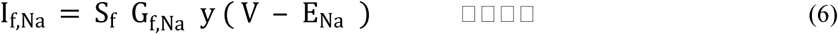

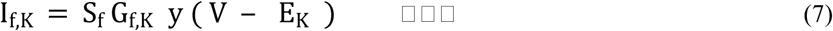

where G_f,Na_ and G_f,K_ are maximal I_f,Na_ and I_f,K_ channel conductance, y is a time-independent inward rectification factor that is a function of voltage, E_Na_, E_K_ are equilibrium potentials of Na^+^ and K^+^ channels respectively, and S_f_ is a scaling factor used to simulate the change of I_f_ expression level.

Formulations of other channel currents for the VM cell model are the same as those in the original model in Ref. (50).

### Evaluating criterion of the pacemaking stability and ability

To analyse the effect of I_K1_ and I_f_ on pacemaking activity, we simulated the membrane potential under different current densities of I_K1_ and I_f_, with I_K1_ being reduced systematically by from 60% to 100% (i.e., I_K1_ density at −80 mV changed from 0.396 to 0 pA/pF while the I_K1_ density in the original basal model is 0.99 pA/pF at −80 mV in I-V curve). The representative I-V relation curve under different inhibition of I_K1_ is shown in S1A Fig. I_f_ density was increased by from 0 to 10 folds with a basal value of −0.63 pA/pF at −80 mV in I-V curve (i.e., I_f_ density changed from 0 to −6.3 pA/pF at −80 mV). The representative I-V relation curve under different incorporation of I_f_ is shown in S1B Fig.

Two characteristics were used to quantify the state of membrane potentials generated by the ventricular pacemaker model: the continuity and validity of spontaneous APs. The continuity was used to quantify whether or not the automaticity of membrane potential could sustain with time; whilst the validity was used to characterize whether every automatic wave was biologically-valid or not. As such, we defined the following:

W: a valid wave. An action potential whose wave trough was less than −20 mV and wave crest was more than 20 mV could be considered as a valid wave.

αW, α<1: an incomplete wave.

R: a resting period lasting 1000 ms.

W^n^: the concatenation of n W’s.

(W R): the concatenation of W and R.

As such, none pacemaking behaviour during the entire simulation period could be described as State-1:

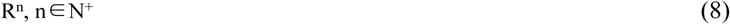

Transient spontaneous pacemaking behaviour could be described as State-2:

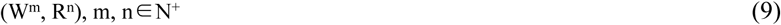

Bursting pacemaking behaviour could be described as State-3:

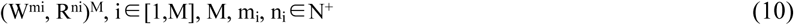

Persistent pacemaking activity with periodically incomplete depolarization could be described as State-4:

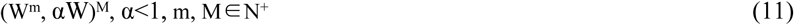

Stable pacemaking activity could be described as State-5:

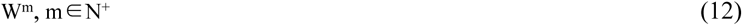

With regard to the pacemaking ability, when the pacemaking behaviour was stable, the cycle length (CL) under varied I_K1_ and I_f_ was calculated. The CL was defined as the averaged wavelength of pacemaking activity over a period of simulation of over 4×10^5^ ms, ensuring the accuracy of the computed CL. As the basal model was for VMs, a long simulation period was necessitated to achieve a completely stable pacemaking status and minimize the effect of the transition period.

### Characteristics of pacemaking during diastolic interval

The length of the diastolic interval (DI) is an important measure to characterize the pacemaking ability. In this study, we defined that DI as the time interval between the time of maximum diastolic potential (MDP) (S6A Fig, t_1_) and the time when the membrane potential reaches at −55 mV (i.e., around the activation potential of the I_CaL_) (S6A Fig, t_2_). The diastolic upstroke velocity during DI was defined as the change rate of the membrane potential, taking the following formulation:

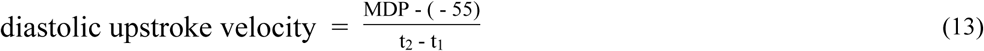

The unit of diastolic upstroke velocity was V/s.

The main inward currents which helped to depolarize membrane potential during DI are I_Na_, I_NaCa_ and I_f_. Their contribution can be described by an average integral during DI:

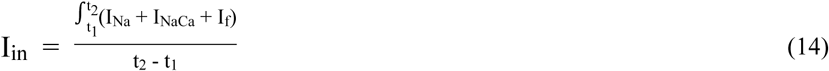

Similarly, the main outward currents which held membrane potential at diastolic potential during DI are I_K1_, I_NaK_, I_Kr_ and I_Ks_, the integral of which can be described as:

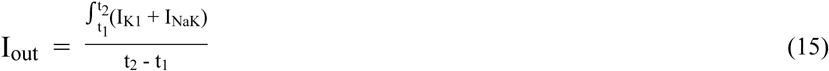

## Supporting Information

**S1 Fig. The I-V curve of I**_**K1**_ **and I**_**f**_ **with different expression level**.

S_K1_ and S_f_ are defined as scaling factors used to simulate the change of I_K1_ and I_f_ expression level. (A) The I-V curve of I_K1_ with S_K1_ of 1, 0.4, 0.1 that gives I_K1_ densities in the I-V curve at −80 mV 0.99, 0.396 and 0.099 pA/pF respectively. (B) The I-V curve of I_f_ with S_f_ of 1, 5, 10 that gives I_f_ densities in the I-V curve at −80 mV −0.63, −3.15 and −6.3 pA/pF respectively.

**S2 Fig. Na**^**+**^**/Ca**^**2+**^ **exchange current (I**_**NaCa**_**) of a transient pacemaking behaviour**.

(A) I_NaCa_ during the entire simulating period of 800 s with the current densities of (I_K1_, I_f_) at (0.297pA/pF, −1.89 pA/pF). (B) Expanded plots of I_NaCa_ traces for the time course from 1.6 ×10^5^ to 1.64 ×10^5^ ms marked by the horizontal brackets with asterisks in (A).

**S3 Fig. Transient spontaneous pacemaking behaviour**.

Membrane potential (V) during the entire simulation period of 400 s with the current densities of (I_K1_, I_f_) at (0.178 pA/pF, −0.63 pA/pF).

**S4 Fig. Na**^**+**^**/Ca**^**2+**^ **exchange current (I**_**NaCa**_**) of a bursting pacemaking behaviour**.

(A) I_NaCa_ with the current densities of (I_K1_, I_f_) at (0.297 pA/pF, −2.52 pA/pF) during the entire simulating period of 800 s. (B) Expanded plots of I_NaCa_ traces for the time course from 3.83×10^5^ ms to 3.85×10^5^ ms marked by the horizontal brackets with asterisks in (A).

**S5 Fig. Partial failure of spontaneous action potentials with different densities of I**_**K1**_.

(A) Membrane potential (V) with the current densities of (I_K1_, I_f_) at (0.297 pA/pF, −3.15 pA/pF) during simulating time course from 3.6×10^5^ to 3.7×10^5^ ms. (B) Membrane potential (V) with the current densities of (I_K1_, I_f_) at (0.277 pA/pF, −3.15 pA/pF) during simulating time course from 3.6×10^5^ to 3.7×10^5^ ms.

**S6 Fig. Change of diastolic depolarizing rate with the increase of I**_**f**_ **density**.

(A) Definition of diastolic depolarizing rate. MDP: maximum diastolic potential; t_1_: the time when membrane potential is MDP; t_2_: the time when potential arrives −55 mV (i.e., around the activation potential of the I_CaL_). (B) Change of diastolic depolarizing rate with the increase of I_f_ density from 0 to −6.3 pA/pF when I_K1_ density at −80 mV is at 0.05 pA/pF.

**S7 Fig. Prolonged cycle length under greater I**_**f**_ **density**.

Membrane potential (V) with the current densities of (I_K1_, I_f_) at (0.05 pA/pF, −3.78 pA/pF) and (0.05 pA/pF, −5.04 pA/pF) (solid and dotted line respectively).

**S8 Fig. Model-dependence test – stable pacemaking behaviour**.

Membrane potential (V) with the current densities of (I_K1_, I_f_) at (0.198 pA/pF, −0.63 pA/pF), (0.178 pA/pF, −0.63 pA/pF) and (0.198 pA/pF, −0.756 pA/pF) (solid, dotted and dashed line respectively) based on a human ventricular myocytes model.

**S9 Fig. Model-dependence test – failed pacemaking activity**.

Membrane potential (V) with the current densities of (I_K1_, I_f_) at (0.198 pA/pF, −1.26 pA/pF) based on a human ventricular myocytes model.

**S1 Text. Model-dependence test**.

## Author Contributions

Conceptualization: Henggui Zhang.

Data curation: Yacong Li.

Formal analysis: Yacong Li, Henggui Zhang, Jules C. Hancox.

Funding acquisition: Henggui Zhang.

Investigation: Yacong Li.

Methodology: Yacong Li, Qince Li, Henggui Zhang.

Project administration: Kuanquan Wang, Henggui Zhang.

Resources: Kuanquan Wang, Henggui Zhang.

Software: Yacong Li, Henggui Zhang.

Supervision: Kuanquan Wang, Henggui Zhang.

Validation: Yacong Li.

Visualization: Yacong Li, Henggui Zhang.

Writing – original draft: Yacong Li.

Writing – review & editing: Henggui Zhang, Jules C. Hancox.

